# Optimized vectors for genetic engineering of *Aureobasidium pullulans*

**DOI:** 10.1101/2025.01.25.634885

**Authors:** Analeigha Colarusso, Audrey Williams, Amy S. Gladfelter, Alison C.E. Wirshing, Daniel J. Lew

## Abstract

*Aureobasidium pullulans* is a polyextremotolerant black yeast that exhibits impressive morphological plasticity. Consequently, it shows promise as a model system for investigating mechanisms of cell adaptation to different environments and the regulation of cell shape. Here, we build upon the current toolkit for working with *A. pullulans* and design and test 25 vectors with seven different codon-optimized fluorophores and three selection cassettes. This includes vectors that allow for dual expression of GFP and mCherry tagged proteins at the *URA3* locus and vectors that enable homology-based deletion or C-terminal tagging of endogenous genes without the need for cloning. This versatile vector series for working with *A. pullulans* will enable a broad range of experiments in this emerging model system.

**SUMMARY:** An optimized plasmid toolkit for genetic engineering of the emerging model fungus, *A. pullulans*.

## INTRODUCTION

The spectacular success of yeast model systems such as *Saccharomyces cerevisiae* and *Schizosaccharomyces pombe* in uncovering the molecular mechanisms underpinning fundamental cell biological processes stems in large part from their experimental tractability. Like the yeast models, many other fungi exhibit convenient features like rapid clonal growth in inexpensive media, efficient homologous recombination enabling facile genome engineering, ability to grow as haploids that facilitate mutant analysis, and resistance to freeze/thaw that allows stable storage. However, only a handful of fungi have well-developed molecular genetic tools like plasmid vectors that facilitate introduction of genetic modifications. Because of differences in promoter behavior and codon usage, such tools must be developed and optimized on a species-by-species basis. Here, we report a suite of plasmids for working with the ubiquitous black yeast *Aureobasidium pullulans*. In combination with efficient chemical transformation (Wirshing et al., 2024), these tools allow for rapid design and construction of *A. pullulans* strains in less than a week without the need for cloning.

*A. pullulans* has recently gained attention for its unconventional multi-budding mode of reproduction (Gostinčar et al., 2014; Mitchison-Field et al., 2019; Petrucco et al., 2024; Wirshing et al., 2024). It also displays an unusual degree of phenotypic plasticity (Slepecky & Starmer 2009) and thrives in many different environments, growing as a polyextremotolerant generalist (Gostinčar and Gunde-Cimerman 2024). This makes *A. pullulans* an excellent model system to ask how cells adapt to different environments, and how growth mode and cell morphology contribute to such adaptation. The new tools reported here make targeted genome modification of *A. pullulans* as easy as it is in other well-established model fungi. They should facilitate use of *A. pullulans* to investigate novel cell biological questions, and will be applicable to further develop its industrial applications, particularly the production of the cell wall polysaccharide pullulan, which is used in the food, pharmaceutical, and cosmetics industries (Gostinčar et al., 2014; Singh et al., 2012).

## RESULTS

In previous work, we showed that some codons in *A. pullulans* could limit expression of commonly used fluorescent reporters, we identified promoters that could be used for heterologous expression, and we developed a straightforward chemical transformation protocol (Wirshing et al., 2024). Here, we build on those findings by developing vectors to enable heterologous expression of fluorescently-tagged proteins, as well as tagging and deletion of endogenous genes.

### Vectors for heterologous expression of fluorescently-tagged proteins

We recently reported the construction of a strain of *A. pullulans* in which the coding region of the endogenous *URA3* gene was replaced with a hygromycin B (Hyg) resistance cassette, making it resistant to Hyg but auxotrophic for uracil (Wirshing et al., 2024) (**Figure 1A**). In that strain background, *URA3*-flanking homology can be used to promote integration of *URA3* and other sequences in place of the Hyg marker, with correct integrants becoming uracil prototrophs and Hyg sensitive. An advantage of this strategy is that it enables integration of desired expression constructs at the *URA3* safe haven site while leaving the strain otherwise wild-type, allowing drug resistance markers to be used for additional genome modifications at other sites.

**Figure 1:**
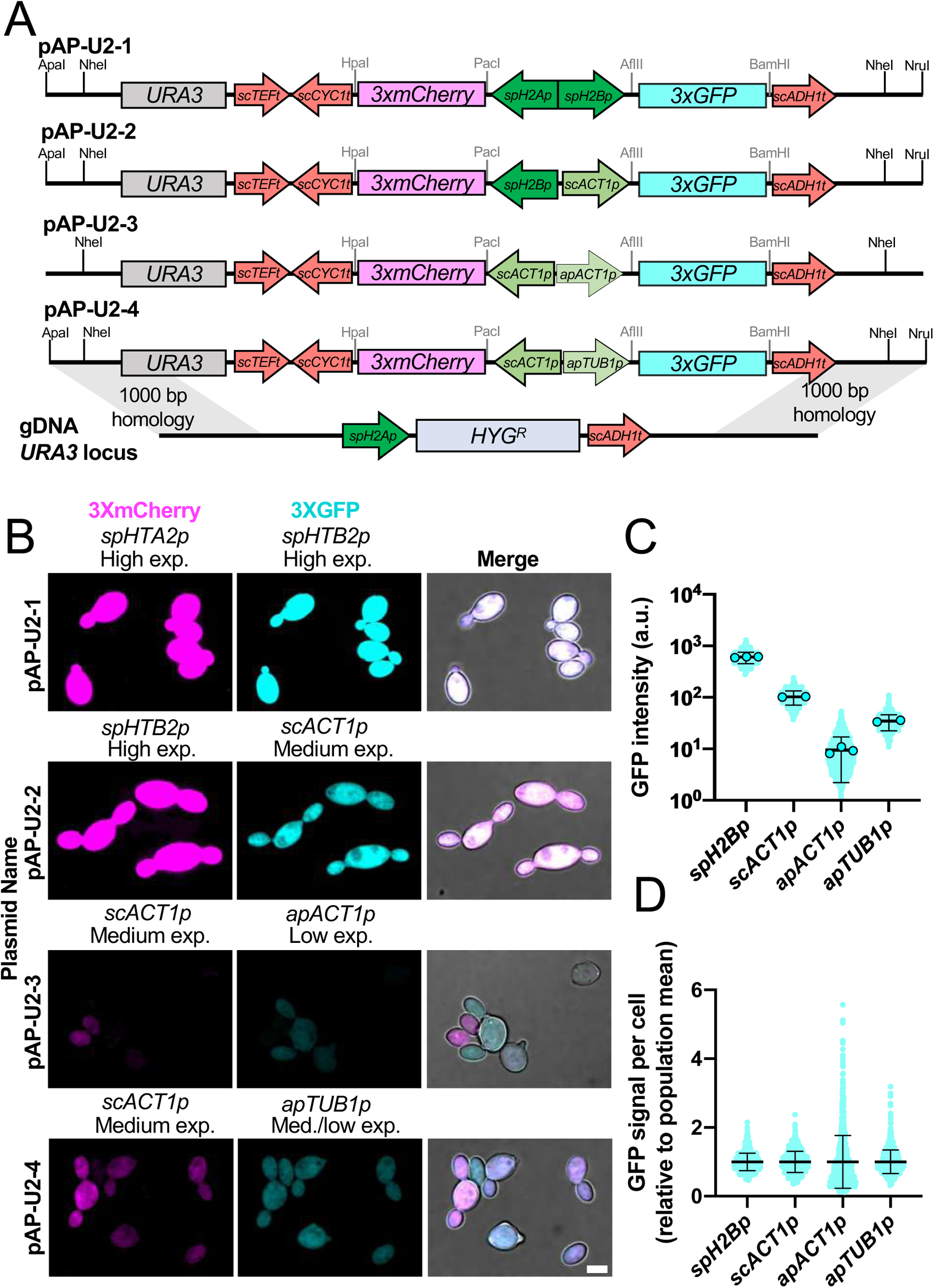
Vectors for dual expression of 3xGFP and 3xmCherry from the native *URA3* locus. (A) Schematic showing the layout of each dual expression cassette, the plasmid name, and the integration strategy to replace *ura3 :: HYG*^*R*^ with the indicated bidirectional expression constructs. Restriction sites for releasing the expression cassette from the plasmid backbone are shown as well as restriction sites for cloning genes adjacent to each fluorophore. Promoters are indicated with “p” and terminators with “t”. (B) Maximum intensity projection of confocal z-stacks of example transformants expressing 3xGFP and 3xmCherry driven by each of the different promoters. All panels displayed with the same intensity scale. Scale bar, 5 µm. (C) Quantification of the average fluorescence intensity in individual cells expressing cytoplasmic 3xGFP under each promoter from two or three separate imaging experiments. Circles represent the mean for each experiment and cyan dots indicate values for individual cells (n = 1328, 896, 1088, and 903 cells) (D) Quantification of the relative GFP intensity per cell relative to the population mean for each promoter tested. This graph highlights differences in cell-to-cell variability across promoters. In both C and D, the mean is shown, and error bars represent the standard deviation.

We had used this strategy to express nuclear and cytoplasmic fluorescent probes from divergent promoters (Wirshing et al., 2024). Here we extend the utility of this approach with four new dual expression vectors (pAP-U2-1 to pAP-U2-4) that similarly use divergent promoters to express three tandem copies of codon-optimized mCherry (3xmCherry) and GFP (3xGFP). The fluorophore coding regions are flanked by convenient restriction sites to allow N- and C-terminal tagging of any desired genes. Each vector has different promoters, with expression levels spanning nearly three orders of magnitude (**Figure 1A-C**). In head-to-head comparisons, the *spH2B* (*Spizellomyces punctatus* histone *H2B*) promoter yielded the highest expression level, followed by the *scACT1* (*S. cerevisiae ACT1*), *apTUB1* (*A. pullulans TUB1*), and *apACT1* (*A. pullulans ACT1*) promoters (**Figure 1B**). This suggests that the *spH2B* promoter is best used when overexpression is desired. Additionally, the *apACT1* promoter results in more variable expression levels than the other promoters tested (**Figure 1D**), and may be most suitable for experiments where variable expression is desired.

pAP-U2 vectors are designed so that unique restriction enzymes (ApaI and NruI, or NheI) can be used to liberate the desired fragment from the vector backbone (**Figure 1A**). Following such digestion, there is sufficient URA3-flanking homology on each side to effectively target integration at the endogenous *URA3* locus (Wirshing et al., 2024). We therefore recommend avoiding the recognition sequences for these restriction enzymes when cloning new genes of interest into these vectors. Digested plasmids can be directly transformed into the uracil auxotrophic strain and transformants are selected on plates lacking uracil. Correct integration at *URA3* (as opposed to integration at other sites in the genome) renders the transformants Hyg sensitive.

### Homology based gene editing without cloning

In addition to heterologous expression, it is frequently desirable to tag or modify endogenous genes. In *S. cerevisiae* or *S. pombe*, where very short homology arms (∼35-80 bp) suffice to target integration by homologous recombination (Bähler et al., 1998a; Baudin et al., 1993; Lorenz et al., 1995; McElver and Weber, 1992), the necessary homology can simply be included in a PCR primer. However, in *A. pullulans* homology arms shorter than 800 bp are significantly less effective than longer homology arms (Wirshing et al., 2024). As this is too long to be included in a primer, we tested whether the homology arms could be amplified separately.

As a test case, we separately amplified 1 kb sequences upstream and downstream of the *URA3* gene, and mixed them with a PCR-amplified HygR cassette with primers designed to include 100 bp of overlap with the homology arms (**Figure 2A**). Transformation with HygR flanked by only 100 bp of homology resulted in very few transformants on Hyg plates, and fewer than 20% of those were uracil auxotrophs (**Figure 2B and 2C**). In contrast, inclusion of the 1 kb flanking homology arms (as separate PCR products) resulted in hundreds of transformants with ∼80% correctly integrated to yield uracil auxotrophs (**Figure 2B and 2C**). Similar results were achieved with only 50 bp of sequence overlap between the homology arms and the selection cassette (**Figure 2D and 2E**). Thus, 3-part-PCR works for targeting integration into the *A. pullulans* genome.

**Figure 2:**
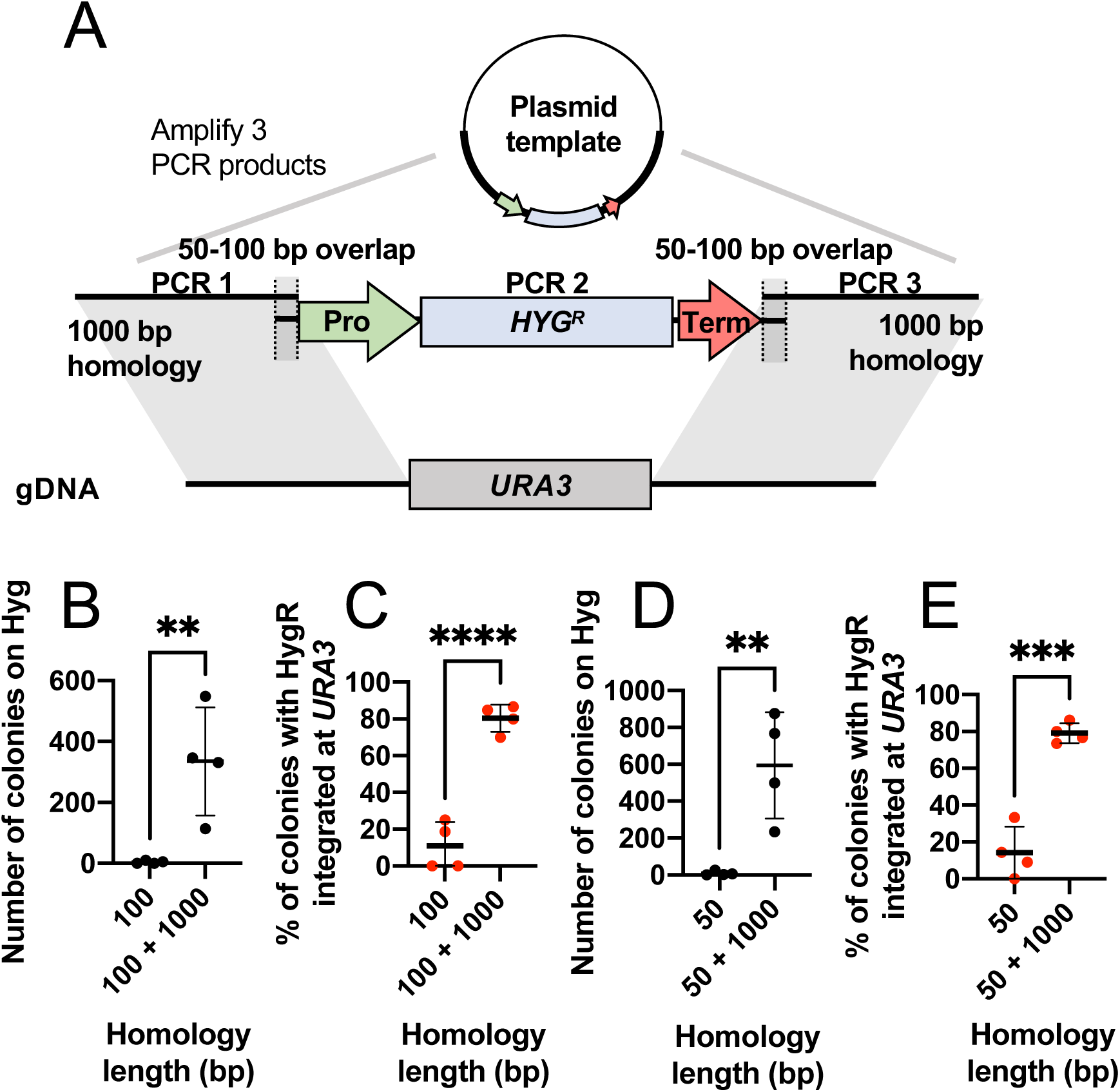
3-part-PCR integration strategy. (A) Schematic showing the strategy to replace *URA3* with a hygromycin-resistance cassette using the 3-part-PCR approach. (B, D) Number of transformants and (C, E) percent of hygromycin-sensitive colonies from four independent transformations with and without 50 bp overlap (B, C) or 100 bp overlap (D, E). Statistical significance calculated by Student’s t test p-values indicated (not shown, p > 0.05; **, p ≤ 0.01; ***, p ≤ 0.001; ****, p ≤ 0.0001).

### Design of PCR-based integration vectors for *A. pullulans*

Based on the success of our 3-part-PCR strategy, we designed a series of PCR-based integration vectors for *A. pullulans* (pAPint) with seven different codon-optimized fluorescent tags and three different selection cassettes (**Figure 3A and 3B**). The selection cassettes confer resistance to Hyg, nourseothricin (Nat), and Geneticin (G418) (Petrucco et al., 2024; Schultz et al., 2019). All resistance genes are driven by the strong *spH2B* promoter with the *scTEF1* terminator and were chosen because they lack the TTA codon that results in poor expression in *A. pullulans* (Petrucco et al., 2024). Minimal inhibitory concentrations (MICs) for Hyg and Nat were reported previously (Petrucco et al., 2024), and the MIC for G418 was 75 µg/ml (**Supplemental Figure 1**). We tested the efficacy of the selection cassettes by using them to replace the native *LEU2* locus (**Figure 3C**). Selection with Hyg, Nat, or G418 yielded hundreds of transformants, with >80% being leucine auxotrophs (**Figure 3D**). Thus, all three cassetts are equally effective.

**Figure 3:**
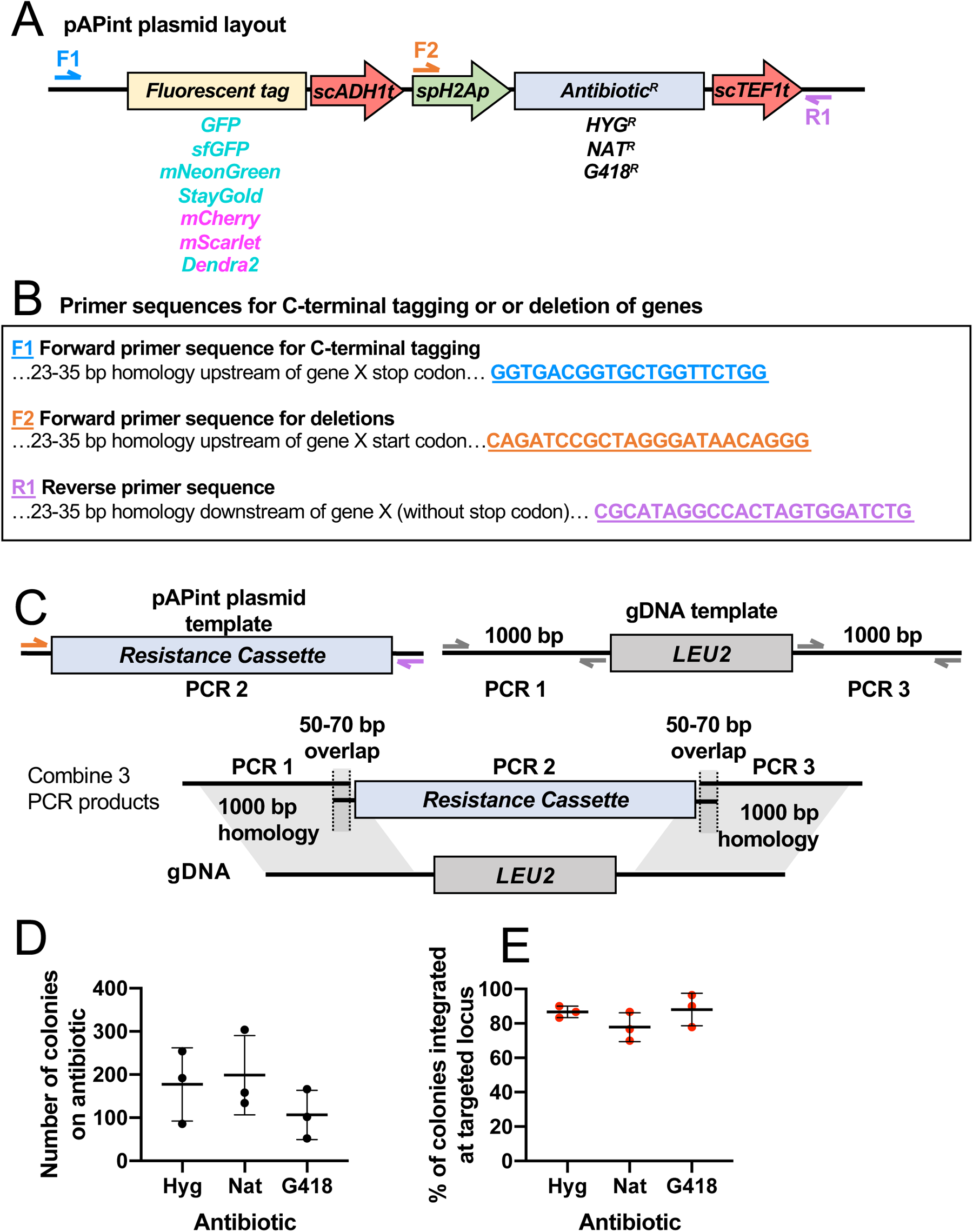
pAPint vectors and primer design. (A) Schematic showing pAPInt vector layout and primer binding sites. (B) Primer sequences for amplifying deletion or C-terminal tagging cassettes from pAPint vectors. (C) Schematic showing strategy to delete *LEU2* and replace it with an antibiotic cassette. (D) Number of transformants and (E) percent integrated at targeted locus for each antibiotic from three independent transformations. In all graphs, the mean and standard deviation are shown. Statistical significance calculated by one-way ANOVA, p > 0.05.

We next used our vectors to C-terminally tag the mitochondrial marker *CIT1* (Wirshing et al., 2024) with the 3-part-PCR method. Correctly integrated transformants (assessed by colony PCR) all exhibited fluorescence localized to mitochondria (**Figure 4A**). Of the red fluorophores, mScarlet was ∼2-fold brighter than mCherry, but ∼7-fold less photostable (**Figure 4B and 4D**). Of the green fluorophores, mNeonGreen (mNG) was about 1.5-fold brighter than GFP, with comparable photostability (**Figure 4C and 4D**). mStayGold (monomeric StayGold) (Ivorra-Molla et al., 2024) was, surprisingly, ∼16-fold dimmer than GFP, but it was ∼1.5-fold more photostable (**Figure 4C and 4D**). The photoconvertible fluorescent protein Dendra2 is green but can be photoconverted to red after exposure to 405 nm or intense 488 nm light (Gurskaya et al., 2006). Before photoconversion, Dendra2 was ∼1.6-fold dimmer than GFP (**Figure 4C**). After photoconversion, Dendra2 was comparable in brightness to mCherry and nearly as photostable (**Figure 4B**). Thus, our pAPint vector series provides multiple different fluorescent tags with different properties to choose from depending on experimental needs.

**Figure 4:**
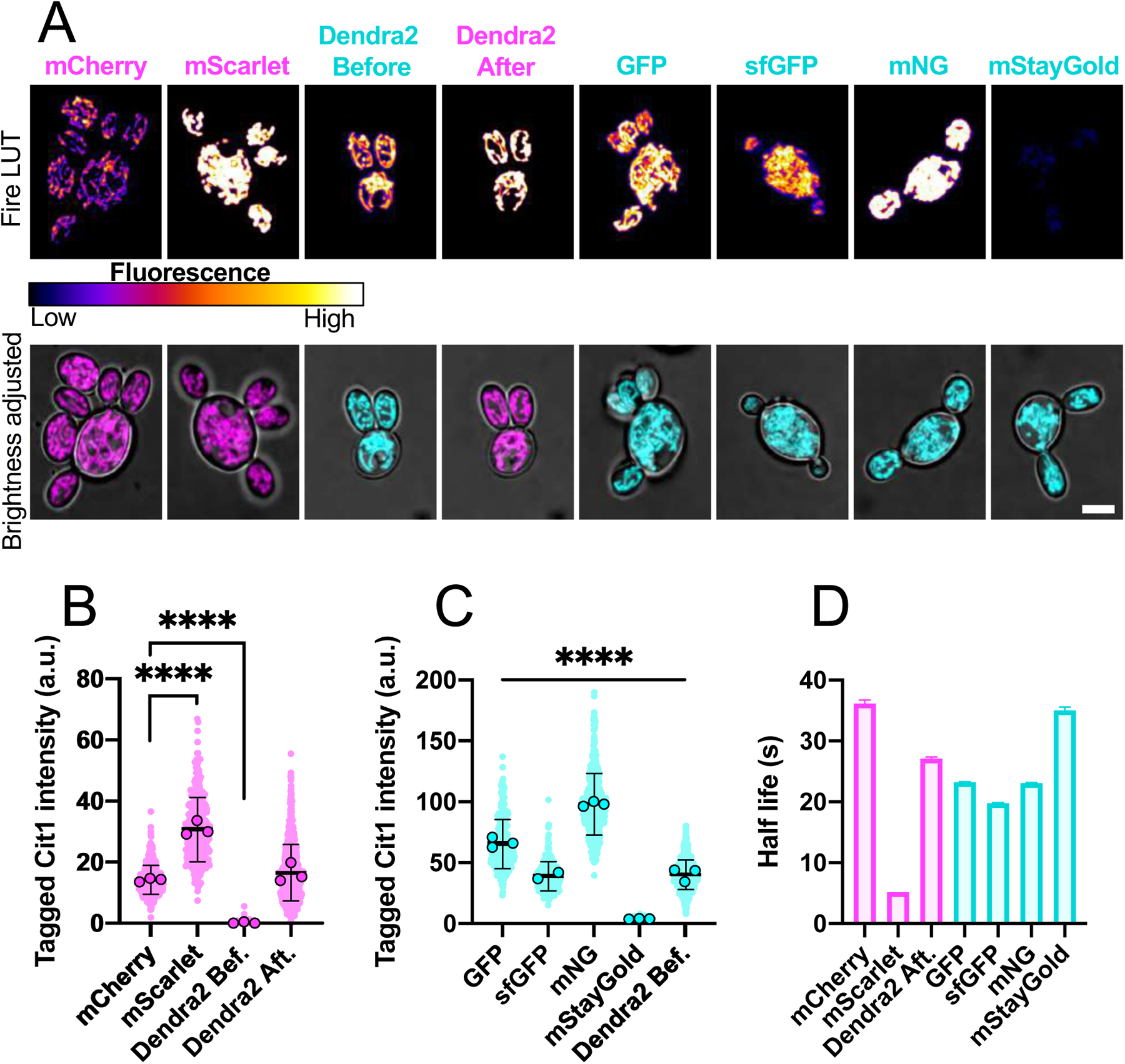
Characterization of codon-optimized fluorophores for C-terminally tagging native genes. (A) Maximum intensity projection of confocal z-stacks showing example transformants expressing Cit1 tagged with each fluorophore. The top row are shown at the same fluoroescense scale in the Fire look up table (LUT) to allow for comparison of intensity across images. In the bottom row, the brightness is adjusted to highlight mitochondrial structure. Scale bar, 5 µm. (B and C) Quantification of the average fluorescence intensity of each cell expressing fluorescently tagged Cit1 from two or three separate imaging experiments. Circles represent the mean for each experiment and dots indicate values for individual cells. Values from cells expressing red fluorescent proteins are shown in B and green fluorescent proteins are shown in C (n = 509 for mCherry, 328 for mScarlet, 226 for GFP, 232 for sfGFP, 431 for mNG, 380 for mStayGold, and 743 for Dendra2). The mean is shown, and error bars represent the standard deviation. (D) Half-life of each fluorophore calculated by fitting the fluorescence decay curves, shown in Supplemental Figure 2, with a one phase decay (n = 15 cells for each condition). Error bars represent the 95% confidence intervals. Statistical significance (B,C) calculated by one way ANOVA, p-values indicated (not shown, p > 0.05; ****, p ≤ 0.0001).

## DISCUSSION

We report an expanded molecular toolkit for working with the emerging model fungus, *A. pullulans*.

The pAP-U2 series vectors enable heterologous expression of 3xGFP and/or 3xmCherry tagged proteins at tunable levels, integrated the *URA3* safe harbor locus. Construction of strains for two-color imaging in a single transformation step is especially useful as traditional sexual crosses are not yet tractable for *A. pullulans* in the lab. Currently, strains carrying multiple mutations (e.g., deletions or fluorescent tags) are generated through successive rounds of transformation to introduce each mutation. As the transformation process is stressful (killing ∼90% of cells) and has the potential to introduce unintended mutations, minimizing transformations is desirable (Wirshing et al., 2024).

We found that highly efficient targeting of homologous recombination to endogenous loci is achieved when flanking sequences are PCR amplified separately from the selection cassette with just 50 bp overlapping sequences between each PCR product. A similar strategy has been used previously to delete a gene in *A. pullulans* (Guo et al., 2017). Based on the efficacy of the 3-part-PCR approach, we designed a series of vectors (pAPint) for PCR-based tagging or deletion of native genes without the need for cloning.

The pAPint series vectors include three different selection cassettes to enable the sequential introduction of multiple different edits in a single strain. We show that *A. pullulans* transformants can be selected using Hyg, Nat, or G418. The pAPint vectors also encode seven different codon-optimized fluorescent proteins for C-terminally tagging of endogenous genes. We included four green (mNG, GFP, sfGFP, and mStayGold) and two red (mScarlet and mCherry) tags with different brightness and photostability characteristics, and one photoswitchable (Dendra2) tag. The fluorescent protein tags were all validated by C-terminal tagging of the mitochondrial marker *CIT1*. In that context, mNG (Shaner et al., 2013) was the brightest green fluorescent protein, and mScarlet (Bindels et al., 2017) the brightest red fluorescent protein. While CIT1-mStayGold (Ivorra-Molla et al., 2024) was the most photostable of the fluorophores tested, it was surprisingly dim. The reason for this is not clear as mStayGold has comparable brightness to mNG in other systems (Hirano et al., 2022). It remains to be determined whether the unexpected dimness of mStayGold reflects some aspect of *A. pullulans* cellular context or is due to the linkage to *CIT1*.

The 25 vectors we present here are analogous to vectors that have been instrumental for working with established model fungi (Bähler et al., 1998b; Longtine et al., 1998; Sheff and Thorn, 2004; Sikorski and Hieter, 1989; Vještica et al., 2020; Wach et al., 1997). This adaptable vector series for genetic engineering in *A. pullulans* should facilitate studies into the fascinating cell biology of this unconventional fungus.

## METHODS

### *A. pullulans* strains and maintenance

Strains used in this study are listed in **Supplemental Table 1**. All experiments were conducted with *Aureobasidium pullulans* strain EXF-150 (Gostinčar et al., 2014). Unless otherwise indicated, *A. pullulans* was grown at 24°C in standard YPD medium (2% glucose, 2% peptone, 1% yeast extract) with 2% BD Bacto^TM^ agar (214050, VWR) in plates.

### Plasmid design

Construction of the backbone used to build the pAP-U2 series of vectors has been described previously (Wirshing et al., 2024). The *S. punctatus* histone H2B and *S. cerevisiae ACT1* promoters were described previously (Petrucco et al. 2024), and we also cloned the native *A. pullulans ACT1* and *TUB1* promoters. The native *ACT1* promoter was defined as 1 kb upstream of the start codon and the *TUB1* promoter as 400 bp upstream of the start codon. Promoters were obtained by PCR from genomic DNA template and assembled into the plasmid backbone using NEBuilder HiFi DNA Assembly Master Mix (E2621L, New England Biolabs).

The pAPint vector series was built using a pFA6A plasmid backbone. These plasmids include different selectable markers under control of the Chytrid *S. punctatus* histone H2B promoter and the *S. cerevisiae ADH1* terminator, previously shown to confer resistance in *A. pullulans* (Petrucco et al., 2024). The HygR gene, *aph(4)-Ia*, and NatR gene, nat, were described previously (Petrucco et al., 2024). The G418 resistance gene, *KanMX*, was amplified from plasmid NM9-SpCas9-NLS3 (Addgene #128177) (Schultz et al., 2019). These antibiotic resistance genes do not have the rare TTA codon that prevents efficient gene expression in *A. pullulans* (Petrucco et al., 2024), and they did not require codon optimization to yield resistance. The sequences for codon-optimized GFP and mCherry were described previously (Wirshing et al., 2024). The sequences for mScarlet, mNG, sfGFP, mStayGold and Dendra2 were codon optimized to eliminate all TTA codons and reduce rare codons (Petrucco et al., 2024), and synthesized by Twist Bioscience (San Francisco, CA, USA) or Integrated DNA Technologies (Coralville, IA, USA). DNA fragments were assembled into the plasmid backbone using NEBuilder HiFi DNA Assembly Master Mix (E2621L, New England Biolabs).

All plasmids were confirmed by Whole Plasmid Sequencing by Plasmidsaurus (Eugene, OR, USA). A list of all plasmids built in this study is in **Supplemental Table 2**. Plasmids and full sequences are available through Addgene.

### Determining appropriate antibiotic selection conditions

A single colony of wild type *A. pullulans* was inoculated into 5 ml YPD and grown overnight at 24°C. Cells were harvested and 10^7^ cells were plated onto YPD plates supplemented with different concentrations of G418 (G64000-5.0, Research Products International) ranging from 0 to 100 µg/ml. Plates were grown for two days at 24°C before counting colonies.

### *A. pullulans* transformation

*A. pullulans* was transformed using PEG/LiAc/ssDNA as described previously (Wirshing et al., 2024). Transformants were selected on YPD plates supplemented with 70.4 µg/ml Hygromycin B (400051-1MJ, Millipore), 75 µg/ml G418 (G64000-5.0, Research Products International), or 50 µg/ml Nourseothricin (N1200-1.0, Research Products International). To select for phototrophs when transforming into our uracil auxotrophic background, cells were selected on media lacking uracil, doURA (6.71 g/L BD Difco^TM^ Yeast Nitrogen Base without Amino Acids, BD291940, FisherScientific, 0.77 g/L Complete Supplement Mixture minus uracil, 1004-100, Sunrise Science Products, 2% glucose, and 2% BD Bacto^TM^ agar, 214050, VWR).

To generate strains expressing 3xGFP and 3xmCherry driven by different bidirectional promoters, plasmids pAPU2-1 through pAP-U2-4 were digested with NheI and directly transformed (without cleanup) into a *ura3::HYG*^*R*^ strain background. Colonies were selected on doURA plates. To identify colonies that integrated the constructs at the targeted *URA3* locus and replaced the HygR cassette, 30 colonies were struck out onto doURA media, grown for 48 hours at 24°C, and then replica plated onto YPD plates supplemented with Hyg. In all cases, >80% of the colonies failed to grow on Hyg. For each construct, three colonies that grew on doURA but not Hyg were screened under the microscope for expression of GFP and mCherry, and all colonies were positive for both fluorophores.

To delete *LEU2* (protein ID 313770) and to tag *CIT1* (protein ID 294441), appropriate primers were used to amplify the upstream and downstream homology arms (using gDNA as a template) and the deletion or tagging cassettes were amplified from different pAPint vectors. A list of all primers used to amplify DNA for transformation is available in **Suppelemntal Table 3**. PCR products were purified and concentrated by adding 0.1 volume of 3 M NaOAc and 2.5 volumes of 100% ethanol followed by incubation on ice for 15 min. The DNA–ethanol mixture was then added to a silica DNA binding column (T1017-2; New England Biolabs), rinsed with 400 μl wash buffer (80% EtOH 20 mM NaCl 2 mM Tris pH 8), allowed to dry for 2.5 min, and then eluted in ∼40 μl 10 mM Tris pH 8. The resulting concentration was ∼1000 ng/µl for each purified PCR product. PCR products were combined at an equimolar ratio (∼5-10 pmol of each) in a final volume of 15-20 µl and added to the cells to be transformed.

To screen transformants for deletion of *LEU2*, 30 individual colonies were struck out onto the appropriate selection media, allowed to grow for two days at 24°C, and replica plated onto media lacking leucine, doLEU (6.71 g/L BD Difco^TM^ Yeast Nitrogen Base without Amino Acids, BD291940, FisherScientific, 0.69 g/L Complete Supplement Mixture minus leucine, 1005-100, Sunrise Science Products, 2% glucose, and 2% BD Bacto^TM^ agar, 214050, VWR). The number of colonies that failed to grow on doLEU was used to calculate the percent that integrated the selection cassette at the targeted *LEU2* locus. To screen for integration at the *CIT1* locus, 3-10 colonies were first screened for expression of fluorescently tagged CIT1 under the microscope. For a subset of colonies correct integration was also confirmed by colony PCR using apCIT1-check-fwd (TCGCAAGTACATCCAGCTAACGC) and ADH1t-check-rev (GCCGGTAGAGGTGTGGTCAATAA) with Phire Plant Direct PCR Master Mix (#F-160S, Thermo Scientific) following the manufacturer’s instructions. All colonies that were positive when screened via microscopy were also positive when screened by colony PCR.

### Live-cell imaging and image analysis

To grow yeast for imaging experiments, a single colony was used to inoculate 5 ml of YPD (2% glucose). Cultures were grown overnight at 24°C to a density of 1-5×10^6^ cells/ml. Cells were pelleted at 9391 rcf for 10 s and resuspended at a final density of ∼7×10^7^ cells/ml. Approximately 2×10^5^ cells were mounted on a 500 µl 1.5% agarose (97062-250, VWR) pad made with CSM (6.71 g/L BD Difco^TM^ Yeast Nitrogen Base without Amino Acids, BD291940, FisherScientific, 0.79 g/L Complete Supplement Mixture, 1001-010, Sunrise Science Products, and 2% glucose). All experiments were conducted at room temperature (20-22°C).

To measure fluorophore brightness, cells were imaged on a Nikon Ti2E inverted microscope with a CSU-W1 spinning-disk head (Yokogawa), CFI60 Plan Apochromat Lambda D 60x Oil Immersion Objective (NA 1.42; Nikon Instruments), and a Hamamatsu ORCA Quest qCMOS camera controlled by NIS-Elements software (Nikon Instruments). The entire cell volume was acquired using 75 Z-slices (at 0.2 μm step intervals). Exposure times of 175 ms at 13.5% laser power (excitation 488 nm) were used to image cells expressing GFP, sfGFP, mNeonGreen, mStayGold, or Dendra2 and exposure times of 175 ms at 20% laser power (excitation 561 nm) were used for cells expressing mCherry, mScarlet, or Dendra2 after photoconversion. To photoconvert Dendra2 the 405 nm laser was used at 100% power and 50 ms exposure. To measure fluorophore photostability, cells were imaged at a single plane at 100 ms intervals for 1.5 minutes with exposure times of 50 ms at 100% laser power for the 488 or 561 lasers depending on the fluorophore being measured.

To quantify the fluorescence intensity of Cit1 tagged with each fluorophore, z-stacks were average intensity projected, and the mean fluorescence intensity of each cell was measured using NIS-Elements General Analysis 3 (GA3, Nikon Instruments) software. Autofluorescence was measured from wild-type cells, not expressing any fluorescent proteins, imaged with the same settings. The average fluorescence signal for each cell was calculated by subtracting the autofluorescence signal from the mean cell signal.

To quantify photostability, the fluorescence intensity over time was measured from 15 cells for each condition using FIJI (Schindelin et al., 2012). The average intensity over time across the 15 cells was plotted in GraphPad Prism and fit with a one phase decay that was used to calculate the fluorescence half-life for each fluorescent protein.

### Statistical analysis

All statistical analysis was done using GraphPad Prism. Unless indicated otherwise, data distributions were assumed to be normal, but this was not formally tested. Statistical comparison between indicated conditions was conducted using the two-sided Student’s t test or one-way ANOVA, as indicated in the figure legends. After running an ANOVA, the Tukey test was used to compare the mean with every other mean. Differences were considered significant if the p-value was <0.05.

## ABBREVIATIONS

DNA: Deoxyribonucleic acid
doLEU: media lacking leucine
doURA: media lacking uracil
G418: Geneticin
Hyg: hygromycin B
MIC: minimal inhibitory concentration
mNG: mNeonGreen
Nat: nourseothricin

## SUPPLEMENTAL MATERIAL

Supplemental Figure 1 shows the results for testing the MIC of G418. Supplemental Figure 2 shows the fluorescence decay over time that was used to calculate the half-life for each fluorophore tested. Supplemental Table 1 provides a list of all strains used in this study. Supplemental Table 2 provides a list of plasmids generated in this study. Supplemental Table 3 provides a list of all primers used to amplify DNA to transform *A. pullulans*.

## DATA AVAILABILITY

The data generated in this study are available from the corresponding author upon reasonable request.

## ACKNOWLEDGEMENTS

We thank Alex Crocker for help with codon optimization, and thanks to the Lew and Gladfelter labs for stimulating discussions. This work was funded by NIH/NIGMS grant R35GM122488 to D.J.L.

## AUTHOR CONTRIBUTIONS

Conceptualization, review, and editing manuscript— A. Colarusso, A.C.E. Wirshing, and D.J. Lew. Data curation, investigation, methodology, visualization, validation, and formal analysis— A. Colarusso, A.C.E. Wirshing and A. Williams. Drafting of manuscript— A. Colarusso and A.C.E. Wirshing. Project administration, supervision, and resources— D.J. Lew and A. S. Gladfelter

## SUPPLEMENTAL FIGURE CAPTIONS

**Supplementary Figure 1:**
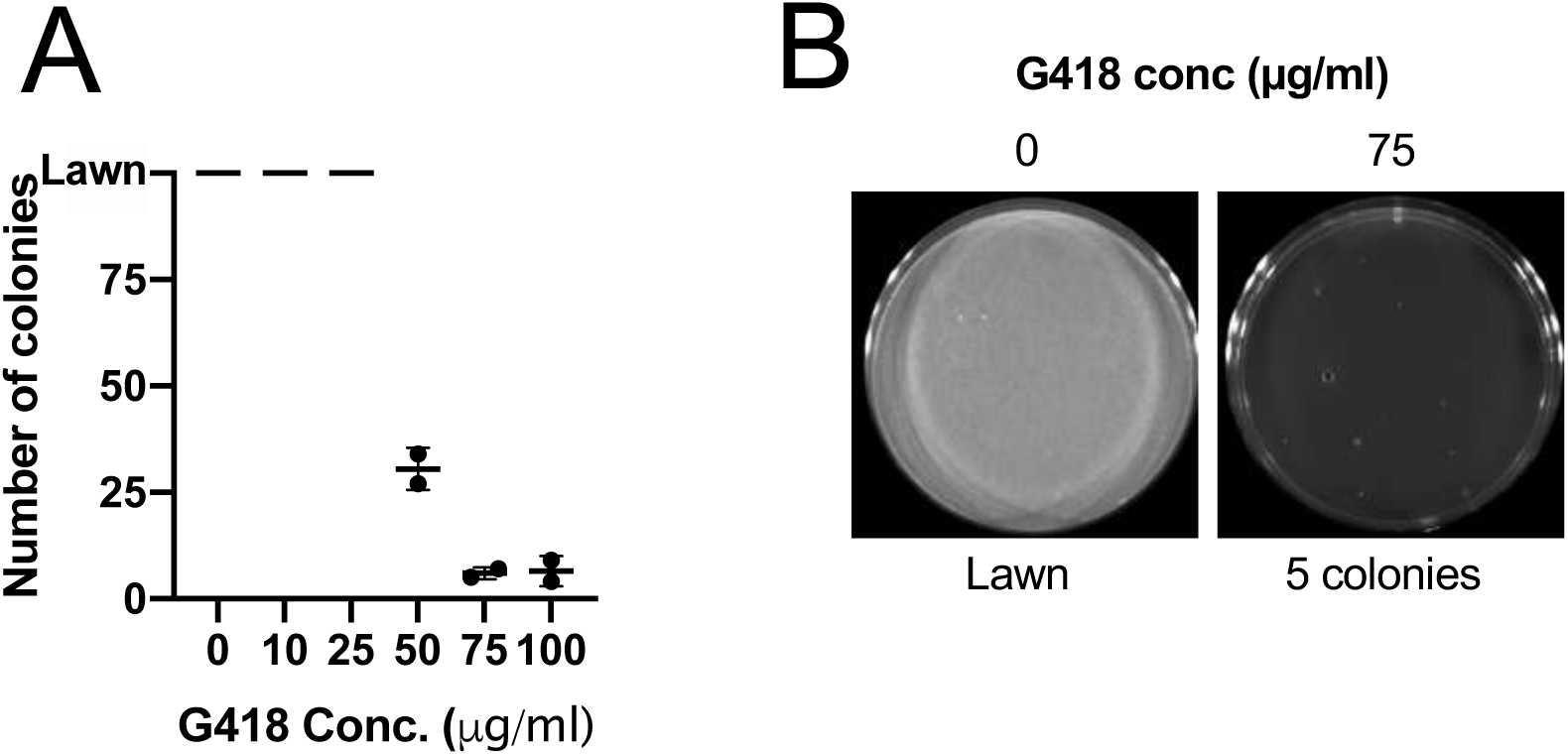
Determination of the minimal inhibitory G418 concentration. (A) Number of colonies present on plates of varying G418 concentrations from two separate experiments. (B) Comparison of plates without G418 and with the concentration used for selection.

**Supplementary Figure 2:**
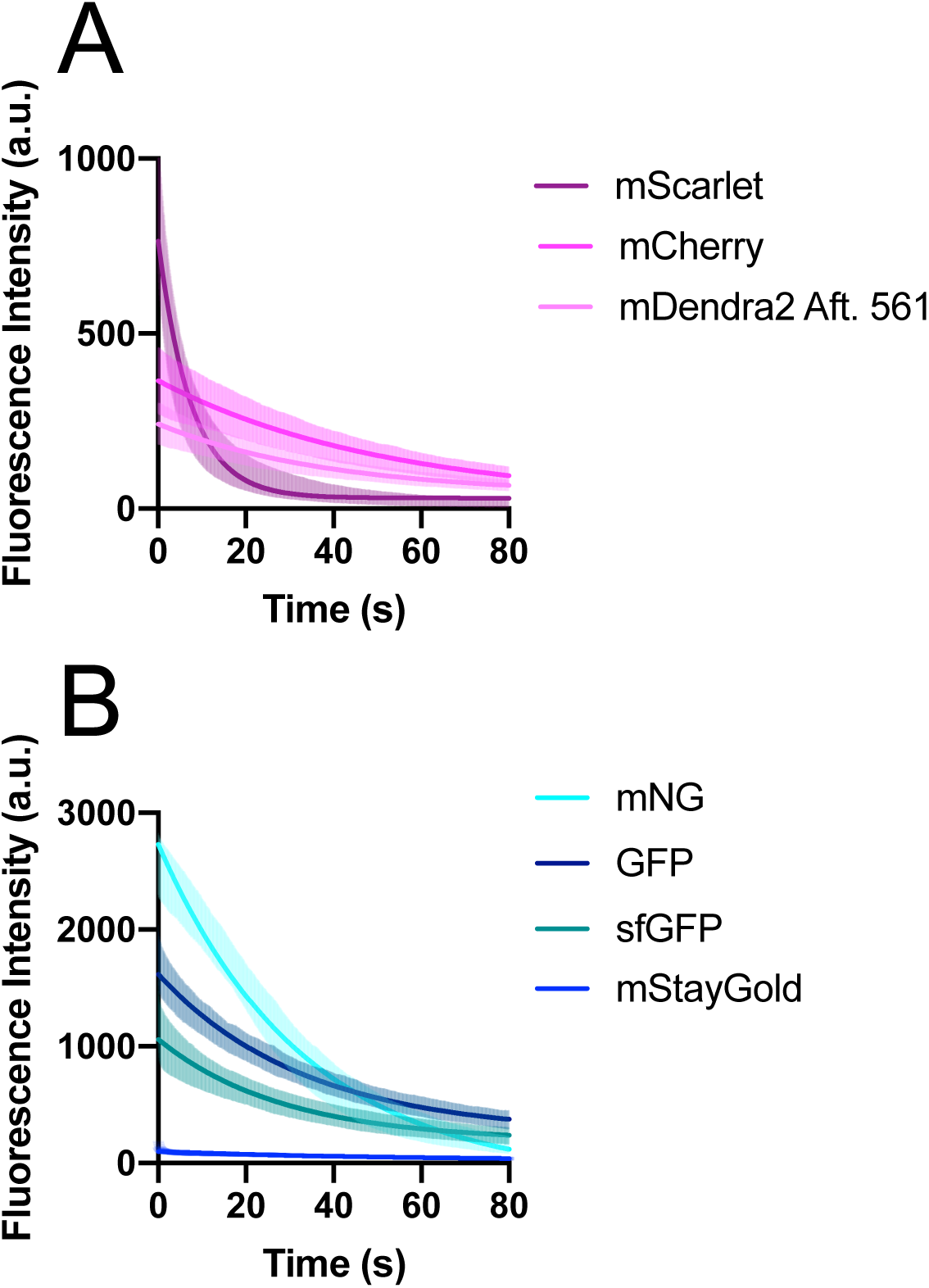
Fluorescence decay curves for each fluorophore. (A-B) Fluorescence decay over time for cells expressing Cit1 tagged with the indicated red fluorescent protein (mScarlet, mCherry, or Dendra2 after photoconversion: A), or green fluorescent protein (mNG, GFP, sfGFP, or mStayGold: B). The mean and standard deviation across 15 cells is shown for each as well as the one phase decay curve fit to each dataset. The R squared values for each curve fitting are 0.75, 0.86, 0.76, 0.92, 0.83, 0.95, 0.78, and 0.90 for mScarlet, mCherry, Dendra2 after, mNG, GFP, sfGFP, and mStayGold, respectively.

**Supplementary Table 1:**
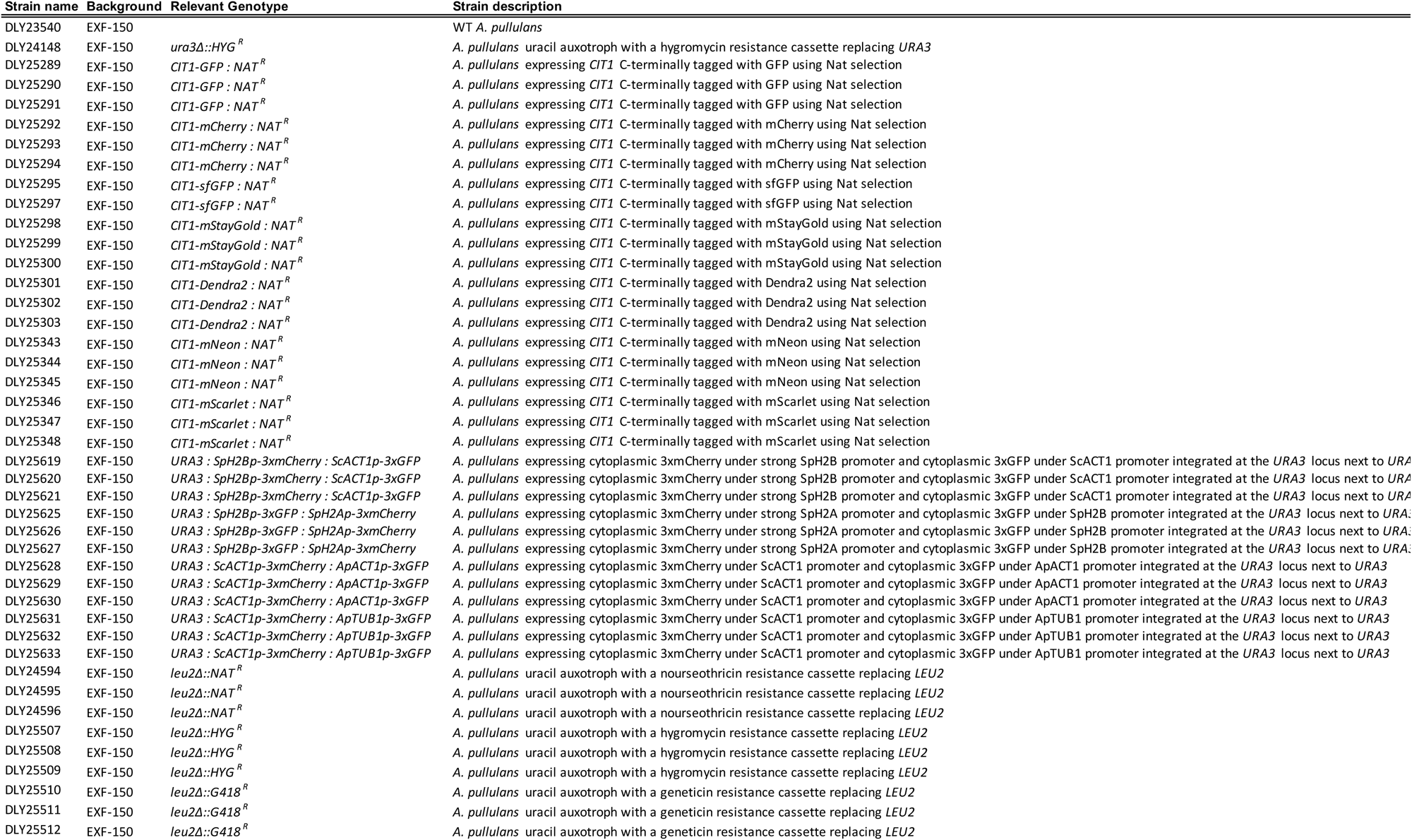

**Supplementary Table 2:**
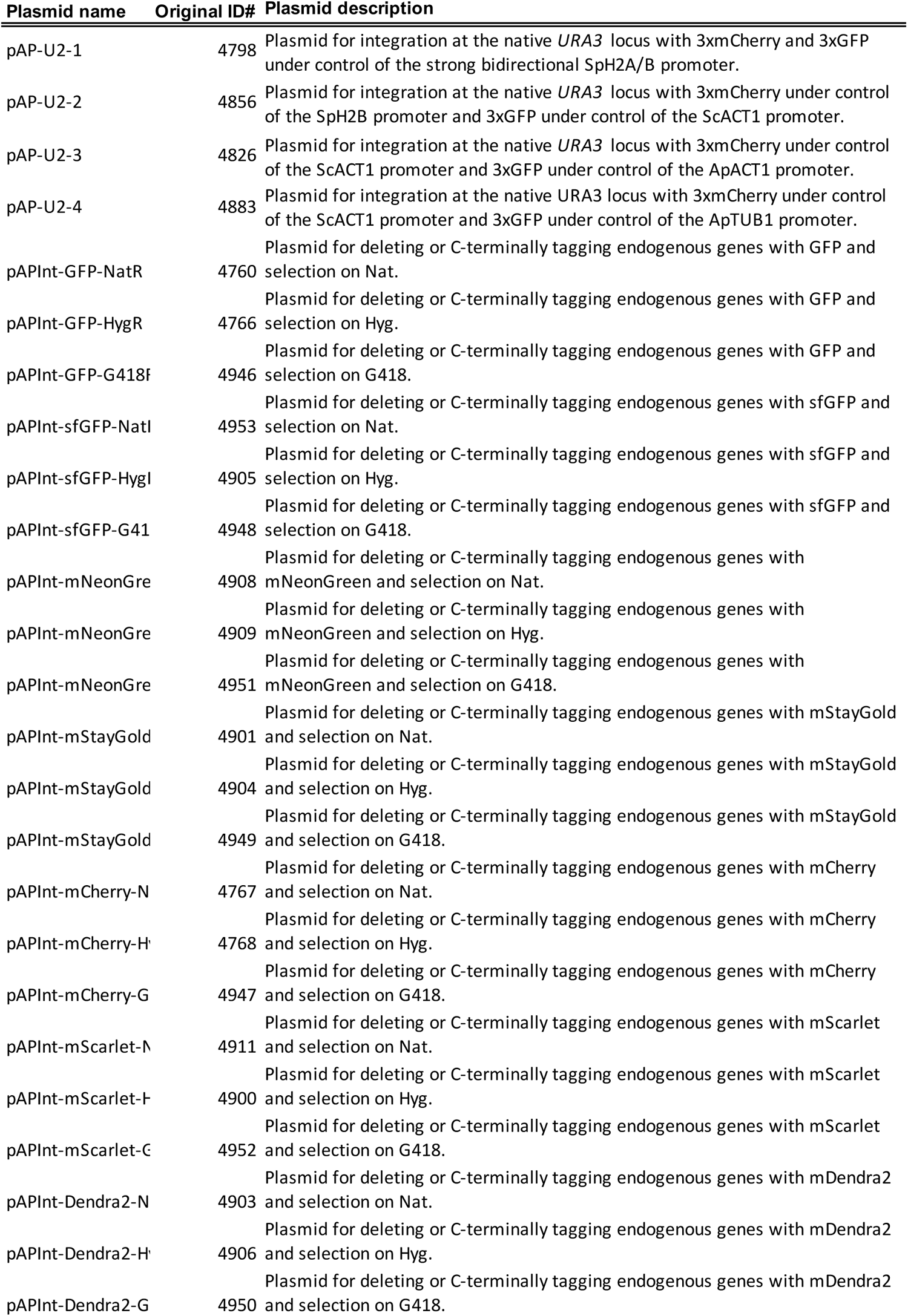

**Supplementary Table 3:**
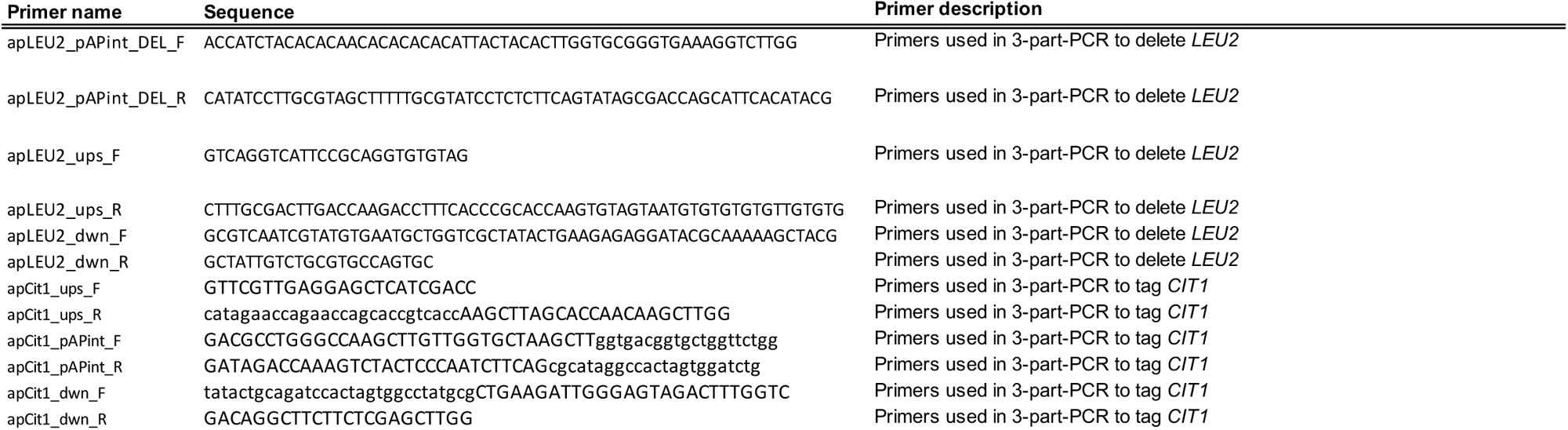

