## Supplemental Table 1 for "Optimized vectors for genetic engineering of *Aureobasidium pullulans*"

| Strain name | Background | Relevant Genotype | Strain description |
|---|---|---|---|
| DLY23540 | EXF-150 | | WT *A. pullulans* |
| DLY24148 | EXF-150 | *ura3Δ::HYG*$^R$ | *A. pullulans* uracil auxotroph with a hygromycin resistance cassette replacing *URA3* |
| DLY25289 | EXF-150 | *CIT1-GFP : NAT*$^R$ | *A. pullulans* expressing *CIT1* C-terminally tagged with GFP using Nat selection |
| DLY25290 | EXF-150 | *CIT1-GFP : NAT*$^R$ | *A. pullulans* expressing *CIT1* C-terminally tagged with GFP using Nat selection |
| DLY25291 | EXF-150 | *CIT1-GFP : NAT*$^R$ | *A. pullulans* expressing *CIT1* C-terminally tagged with GFP using Nat selection |
| DLY25292 | EXF-150 | *CIT1-mCherry : NAT*$^R$ | *A. pullulans* expressing *CIT1* C-terminally tagged with mCherry using Nat selection |
| DLY25293 | EXF-150 | *CIT1-mCherry : NAT*$^R$ | *A. pullulans* expressing *CIT1* C-terminally tagged with mCherry using Nat selection |
| DLY25294 | EXF-150 | *CIT1-mCherry : NAT*$^R$ | *A. pullulans* expressing *CIT1* C-terminally tagged with mCherry using Nat selection |
| DLY25295 | EXF-150 | *CIT1-sfGFP : NAT*$^R$ | *A. pullulans* expressing *CIT1* C-terminally tagged with sfGFP using Nat selection |
| DLY25297 | EXF-150 | *CIT1-sfGFP : NAT*$^R$ | *A. pullulans* expressing *CIT1* C-terminally tagged with sfGFP using Nat selection |
| DLY25298 | EXF-150 | *CIT1-mStayGold : NAT*$^R$ | *A. pullulans* expressing *CIT1* C-terminally tagged with mStayGold using Nat selection |
| DLY25299 | EXF-150 | *CIT1-mStayGold : NAT*$^R$ | *A. pullulans* expressing *CIT1* C-terminally tagged with mStayGold using Nat selection |
| DLY25300 | EXF-150 | *CIT1-mStayGold : NAT*$^R$ | *A. pullulans* expressing *CIT1* C-terminally tagged with mStayGold using Nat selection |
| DLY25301 | EXF-150 | *CIT1-Dendra2 : NAT*$^R$ | *A. pullulans* expressing *CIT1* C-terminally tagged with Dendra2 using Nat selection |
| DLY25302 | EXF-150 | *CIT1-Dendra2 : NAT*$^R$ | *A. pullulans* expressing *CIT1* C-terminally tagged with Dendra2 using Nat selection |
| DLY25303 | EXF-150 | *CIT1-Dendra2 : NAT*$^R$ | *A. pullulans* expressing *CIT1* C-terminally tagged with Dendra2 using Nat selection |
| DLY25343 | EXF-150 | *CIT1-mNeon : NAT*$^R$ | *A. pullulans* expressing *CIT1* C-terminally tagged with mNeon using Nat selection |
| DLY25344 | EXF-150 | *CIT1-mNeon : NAT*$^R$ | *A. pullulans* expressing *CIT1* C-terminally tagged with mNeon using Nat selection |
| DLY25345 | EXF-150 | *CIT1-mNeon : NAT*$^R$ | *A. pullulans* expressing *CIT1* C-terminally tagged with mNeon using Nat selection |
| DLY25346 | EXF-150 | *CIT1-mScarlet : NAT*$^R$ | *A. pullulans* expressing *CIT1* C-terminally tagged with mScarlet using Nat selection |
| DLY25347 | EXF-150 | *CIT1-mScarlet : NAT*$^R$ | *A. pullulans* expressing *CIT1* C-terminally tagged with mScarlet using Nat selection |
| DLY25348 | EXF-150 | *CIT1-mScarlet : NAT*$^R$ | *A. pullulans* expressing *CIT1* C-terminally tagged with mScarlet using Nat selection |
| DLY25619 | EXF-150 | *URA3 : SpH2Bp-3xmCherry : ScACT1p-3xGFP* | *A. pullulans* expressing cytoplasmic 3xmCherry under strong SpH2B promoter and cytoplasmic 3xGFP under ScACT1 promoter integrated at the *URA3* locus next to *URA* |
| DLY25620 | EXF-150 | *URA3 : SpH2Bp-3xmCherry : ScACT1p-3xGFP* | *A. pullulans* expressing cytoplasmic 3xmCherry under strong SpH2B promoter and cytoplasmic 3xGFP under ScACT1 promoter integrated at the *URA3* locus next to *URA* |
| DLY25621 | EXF-150 | *URA3 : SpH2Bp-3xmCherry : ScACT1p-3xGFP* | *A. pullulans* expressing cytoplasmic 3xmCherry under strong SpH2B promoter and cytoplasmic 3xGFP under ScACT1 promoter integrated at the *URA3* locus next to *URA* |
| DLY25625 | EXF-150 | *URA3 : SpH2Bp-3xGFP : SpH2Ap-3xmCherry* | *A. pullulans* expressing cytoplasmic 3xmCherry under strong SpH2A promoter and cytoplasmic 3xGFP under SpH2B promoter integrated at the *URA3* locus next to *URA3* |
| DLY25626 | EXF-150 | *URA3 : SpH2Bp-3xGFP : SpH2Ap-3xmCherry* | *A. pullulans* expressing cytoplasmic 3xmCherry under strong SpH2A promoter and cytoplasmic 3xGFP under SpH2B promoter integrated at the *URA3* locus next to *URA3* |
| DLY25627 | EXF-150 | *URA3 : SpH2Bp-3xGFP : SpH2Ap-3xmCherry* | *A. pullulans* expressing cytoplasmic 3xmCherry under strong SpH2A promoter and cytoplasmic 3xGFP under SpH2B promoter integrated at the *URA3* locus next to *URA3* |
| DLY25628 | EXF-150 | *URA3 : ScACT1p-3xmCherry : ApACT1p-3xGFP* | *A. pullulans* expressing cytoplasmic 3xmCherry under ScACT1 promoter and cytoplasmic 3xGFP under ApACT1 promoter integrated at the *URA3* locus next to *URA3* |
| DLY25629 | EXF-150 | *URA3 : ScACT1p-3xmCherry : ApACT1p-3xGFP* | *A. pullulans* expressing cytoplasmic 3xmCherry under ScACT1 promoter and cytoplasmic 3xGFP under ApACT1 promoter integrated at the *URA3* locus next to *URA3* |
| DLY25630 | EXF-150 | *URA3 : ScACT1p-3xmCherry : ApACT1p-3xGFP* | *A. pullulans* expressing cytoplasmic 3xmCherry under ScACT1 promoter and cytoplasmic 3xGFP under ApACT1 promoter integrated at the *URA3* locus next to *URA3* |
| DLY25631 | EXF-150 | *URA3 : ScACT1p-3xmCherry : ApTUB1p-3xGFP* | *A. pullulans* expressing cytoplasmic 3xmCherry under ScACT1 promoter and cytoplasmic 3xGFP under ApTUB1 promoter integrated at the *URA3* locus next to *URA3* |
| DLY25632 | EXF-150 | *URA3 : ScACT1p-3xmCherry : ApTUB1p-3xGFP* | *A. pullulans* expressing cytoplasmic 3xmCherry under ScACT1 promoter and cytoplasmic 3xGFP under ApTUB1 promoter integrated at the *URA3* locus next to *URA3* |
| DLY25633 | EXF-150 | *URA3 : ScACT1p-3xmCherry : ApTUB1p-3xGFP* | *A. pullulans* expressing cytoplasmic 3xmCherry under ScACT1 promoter and cytoplasmic 3xGFP under ApTUB1 promoter integrated at the *URA3* locus next to *URA3* |
| DLY24594 | EXF-150 | *leu2Δ::NAT*$^R$ | *A. pullulans* uracil auxotroph with a nourseothricin resistance cassette replacing *LEU2* |
| DLY24595 | EXF-150 | *leu2Δ::NAT*$^R$ | *A. pullulans* uracil auxotroph with a nourseothricin resistance cassette replacing *LEU2* |
| DLY24596 | EXF-150 | *leu2Δ::NAT*$^R$ | *A. pullulans* uracil auxotroph with a nourseothricin resistance cassette replacing *LEU2* |
| DLY25507 | EXF-150 | *leu2Δ::HYG*$^R$ | *A. pullulans* uracil auxotroph with a hygromycin resistance cassette replacing *LEU2* |
| DLY25508 | EXF-150 | *leu2Δ::HYG*$^R$ | *A. pullulans* uracil auxotroph with a hygromycin resistance cassette replacing *LEU2* |
| DLY25509 | EXF-150 | *leu2Δ::HYG*$^R$ | *A. pullulans* uracil auxotroph with a hygromycin resistance cassette replacing *LEU2* |
| DLY25510 | EXF-150 | *leu2Δ::G418*$^R$ | *A. pullulans* uracil auxotroph with a geneticin resistance cassette replacing *LEU2* |
| DLY25511 | EXF-150 | *leu2Δ::G418*$^R$ | *A. pullulans* uracil auxotroph with a geneticin resistance cassette replacing *LEU2* |
| DLY25512 | EXF-150 | *leu2Δ::G418*$^R$ | *A. pullulans* uracil auxotroph with a geneticin resistance cassette replacing *LEU2* |
