## Supplemental Table 2 for "Optimized vectors for genetic engineering of *Aureobasidium pullulans*"

| Plasmid name | Original ID# | Plasmid description |
|---|---|---|
| pAP-U2-1 | 4798 | Plasmid for integration at the native *URA3* locus with 3xmCherry and 3xGFP under control of the strong bidirectional SpH2A/B promoter. |
| pAP-U2-2 | 4856 | Plasmid for integration at the native *URA3* locus with 3xmCherry under control of the SpH2B promoter and 3xGFP under control of the ScACT1 promoter. |
| pAP-U2-3 | 4826 | Plasmid for integration at the native *URA3* locus with 3xmCherry under control of the ScACT1 promoter and 3xGFP under control of the ApACT1 promoter. |
| pAP-U2-4 | 4883 | Plasmid for integration at the native URA3 locus with 3xmCherry under control of the ScACT1 promoter and 3xGFP under control of the ApTUB1 promoter. |
| pAPInt-GFP-NatR | 4760 | Plasmid for deleting or C-terminally tagging endogenous genes with GFP and selection on Nat. |
| pAPInt-GFP-HygR | 4766 | Plasmid for deleting or C-terminally tagging endogenous genes with GFP and selection on Hyg. |
| pAPInt-GFP-G418F | 4946 | Plasmid for deleting or C-terminally tagging endogenous genes with GFP and selection on G418. |
| pAPInt-sfGFP-NatI | 4953 | Plasmid for deleting or C-terminally tagging endogenous genes with sfGFP and selection on Nat. |
| pAPInt-sfGFP-HygI | 4905 | Plasmid for deleting or C-terminally tagging endogenous genes with sfGFP and selection on Hyg. |
| pAPInt-sfGFP-G41 | 4948 | Plasmid for deleting or C-terminally tagging endogenous genes with sfGFP and selection on G418. |
| pAPInt-mNeonGre | 4908 | Plasmid for deleting or C-terminally tagging endogenous genes with mNeonGreen and selection on Nat. |
| pAPInt-mNeonGre | 4909 | Plasmid for deleting or C-terminally tagging endogenous genes with mNeonGreen and selection on Hyg. |
| pAPInt-mNeonGre | 4951 | Plasmid for deleting or C-terminally tagging endogenous genes with mNeonGreen and selection on G418. |
| pAPInt-mStayGold | 4901 | Plasmid for deleting or C-terminally tagging endogenous genes with mStayGold and selection on Nat. |
| pAPInt-mStayGold | 4904 | Plasmid for deleting or C-terminally tagging endogenous genes with mStayGold and selection on Hyg. |
| pAPInt-mStayGold | 4949 | Plasmid for deleting or C-terminally tagging endogenous genes with mStayGold and selection on G418. |
| pAPInt-mCherry-N | 4767 | Plasmid for deleting or C-terminally tagging endogenous genes with mCherry and selection on Nat. |
| pAPInt-mCherry-H | 4768 | Plasmid for deleting or C-terminally tagging endogenous genes with mCherry and selection on Hyg. |
| pAPInt-mCherry-G | 4947 | Plasmid for deleting or C-terminally tagging endogenous genes with mCherry and selection on G418. |
| pAPInt-mScarlet-N | 4911 | Plasmid for deleting or C-terminally tagging endogenous genes with mScarlet and selection on Nat. |
| pAPInt-mScarlet-H | 4900 | Plasmid for deleting or C-terminally tagging endogenous genes with mScarlet and selection on Hyg. |
| pAPInt-mScarlet-G | 4952 | Plasmid for deleting or C-terminally tagging endogenous genes with mScarlet and selection on G418. |
| pAPInt-Dendra2-N | 4903 | Plasmid for deleting or C-terminally tagging endogenous genes with mDendra2 and selection on Nat. |
| pAPInt-Dendra2-H | 4906 | Plasmid for deleting or C-terminally tagging endogenous genes with mDendra2 and selection on Hyg. |
| pAPInt-Dendra2-G | 4950 | Plasmid for deleting or C-terminally tagging endogenous genes with mDendra2 and selection on G418. |
