## Supplemental Table 3 for "Optimized vectors for genetic engineering of *Aureobasidium pullulans*"

| Primer name | Sequence | Primer description |
|---|---|---|
| apLEU2_pAPint_DEL_F | ACCATCTACACACAACACACACACATTACTACACTTGGTGCGGGTGAAAGGTCTTGG | Primers used in 3-part-PCR to delete *LEU2* |
| apLEU2_pAPint_DEL_R | CATATCCTTGCGTAGCTTTTTGCGTATCCTCTCTTCAGTATAGCGACCAGCATTCACATACG | Primers used in 3-part-PCR to delete *LEU2* |
| apLEU2_ups_F | GTCAGGTCATTCCGCAGGTGTGTAG | Primers used in 3-part-PCR to delete *LEU2* |
| apLEU2_ups_R | CTTTGCGACTTGACCAAGACCTTTCACCCGCACCAAGTGTAGTAATGTGTGTGTGTTGTGTG | Primers used in 3-part-PCR to delete *LEU2* |
| apLEU2_dwn_F | GCGTCAATCGTATGTGAATGCTGGTCGCTATACTGAAGAGAGGATACGCAAAAAGCTACG | Primers used in 3-part-PCR to delete *LEU2* |
| apLEU2_dwn_R | GCTATTGTCTGCGTGCCAGTGC | Primers used in 3-part-PCR to delete *LEU2* |
| apCit1_ups_F | GTTCGTTGAGGAGCTCATCGACC | Primers used in 3-part-PCR to tag *CIT1* |
| apCit1_ups_R | catagaaccagaaccagcaccgtcaccAAGCTTAGCACCAACAAGCTTGG | Primers used in 3-part-PCR to tag *CIT1* |
| apCit1_pAPint_F | GACGCCTGGGCCAAGCTTGTTGGTGCTAAGCTTggtgacggtgctggttctgg | Primers used in 3-part-PCR to tag *CIT1* |
| apCit1_pAPint_R | GATAGACCAAAGTCTACTCCCAATCTTCAGcgcataggccactagtggatctg | Primers used in 3-part-PCR to tag *CIT1* |
| apCit1_dwn_F | tatactgcagatccactagtggcctatgcgCTGAAGATTGGGAGTAGACTTTGGTC | Primers used in 3-part-PCR to tag *CIT1* |
| apCit1_dwn_R | GACAGGCTTCTTCTCGAGCTTGG | Primers used in 3-part-PCR to tag *CIT1* |
